# BOston Neonatal Brain Injury Dataset for Hypoxic Ischemic Encephalopathy (BONBID-HIE): Part I. MRI and Manual Lesion Annotation

**DOI:** 10.1101/2023.06.30.546841

**Authors:** Rina Bao, Ya’nan Song, Sara V. Bates, Rebecca J. Weiss, Anna N. Foster, Camilo Jaimes Cobos, Susan Sotardi, Yue Zhang, Randy L. Gollub, P. Ellen Grant, Yangming Ou

## Abstract

Hypoxic ischemic encephalopathy (HIE) is a brain injury that occurs in 1 ∼5*/*1000 term neonates. Accurate identification and segmentation of HIE-related lesions in neonatal brain magnetic resonance images (MRIs) is the first step toward predicting prognosis, identifying high-risk patients, and evaluating treatment effects. It will lead to a more accurate estimation of prognosis, a better understanding of neurological symptoms, and a timely prediction of response to therapy. We release the first public dataset containing neonatal brain diffusion MRI and expert annotation of lesions from 133 patients diagnosed with HIE. HIE-related lesions in brain MRI are often diffuse (i.e., multi-focal), and small (over half the patients in our data having lesions occupying <1% of brain volume). Segmentation for HIE MRI data is remarkably different from, and arguably more challenging than, other segmentation tasks such as brain tumors with focal and relatively large lesions. We hope that this dataset can help fuel the development of MRI lesion segmentation methods for HIE and small diffuse lesions in general.

## Background & Summary

Accurate identification of brain lesion injuries in neonatal brain magnetic resonance images (MRI) [1, 2, 3] is crucial to improve clinical care of neonates with hypoxic ischemic encephalopathy (HIE), a brain disease that occurs in around 1 ∼5*/*1000 term-born infants at birth [4, 5]. HIE affects around 200,000 term-born neonates every year worldwide [4, 5], costing about $2 billion/year in the US alone, let alone family burdens. Therapeutic hypothermia, the current clinical treatment of HIE, can reduce mortality and morbidity. Nevertheless, around 60% of patients still die or develop neurocognitive deficits by 2 years of age. MRI is used in over 50% of all the >100 ongoing HIE-related clinical trials worldwide [6], for evaluating treatment effects [7, 8, 9], and helping discover clinical [10, 11, 12], biochemical [10, 13, 14, 15], and serum [16, 17, 18] biomarkers. Accurate identification of brain lesions in neonatal brain MRIs [1, 2, 3] is needed for disease prognosis, a better understanding of the neural basis of disease progression, and more timely evaluations of therapeutic effects.

HIE lesions are often diffuse (i.e., multi-focal), and small; hence, algorithms that have shown great promise in segmenting big and focal lesions, such as brain tumors and acute strokes, often encounter challenges when directly applied to MRIs of HIE patients. Indeed, many (over half) patients had lesions occupying <1% of brain volume, as shown in Figure 1. As a result, the segmentation accuracy measured by the Dice overlap with U-Net [19] and other state-of-the-art machine/deep learning algorithms on HIE remains at around 0.5 [20], whereas Dice is over 0.8 when segmenting brain tumors [21, 22].

**Figure 1.**
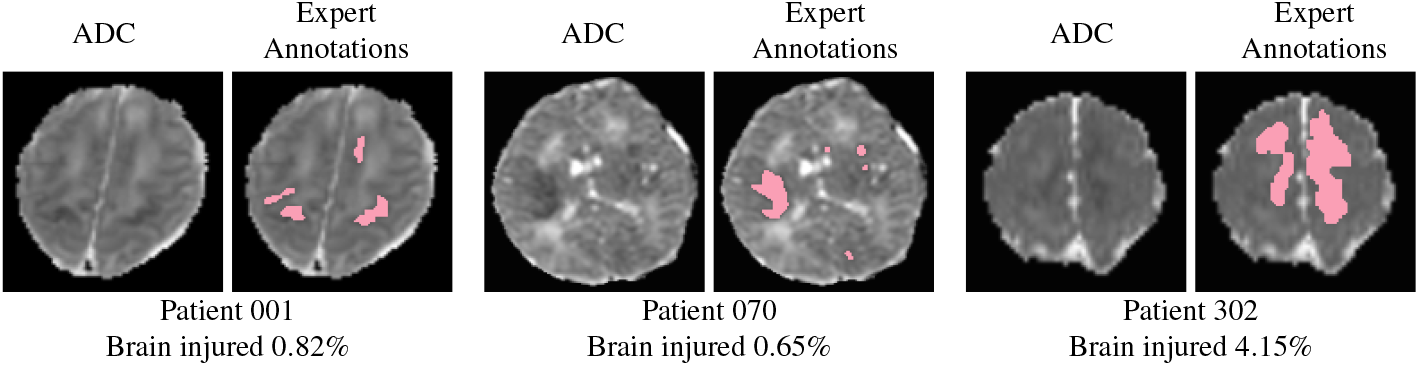
Lesions associated to hypoxic ischemic encephalopathy (HIE) are typically diffuse (i.e., multi-focal) and small. Here we show two representative images for 3 HIE patients. For each patient, in the left panel: apparent diffusion coefficient (ADC) maps that are clinically used to identify HIE lesions; in the right panel: manually-annotated lesions (shown in pink) overlaid on the ADC map. We listed the percentage of the whole brain volume being injured at the bottom (i.e., lesion volume divided by the whole brain volume).

A major hurdle in developing algorithms for small diffuse lesions, such as HIE lesions, is the lack of public data. Public data with expert annotations of lesions have fueled the advancement of machine learning algorithms to segment brain tumors [23], stroke lesions [24], multiple sclerosis lesions [25, 26], and numerous other diseases in the brain or other organs [27, 28]. However, to date, there is no public MRI data with expert annotations available for HIE lesions.

We present BOston Neonatal Brain Injury Dataset for Hypoxic Ischemic Encephalopathy (BONBID-HIE), an open-source, comprehensive, and representative MRI dataset for HIE. This paper introduces the first part of the BONBID-HIE data. This release contains raw and derived diffusion parameter maps, as well as manually-annotated lesion masks, for 133 HIE patients.

Our data was from Massachusetts General Hospital. It includes MRIs from different scanners (Siemens 3T and GE 1.5T), different MRI protocols, and from patients of different races/ethnicities and ages (0-14 days postnatal age). Part I of our data release (this paper) focuses on lesion detection, while Part II (a follow-up paper) will focus on clinical, treatment, and neurologic outcome data for further developing prognostic biomarkers.

## Methods

This work was approved by the Institutional Review Boards (IRBs) at Massachusetts General Hospital (MGH) and Boston Children’s Hospital (BCH). Figure 2 illustrates the overall data archiving process.

**Figure 2.**
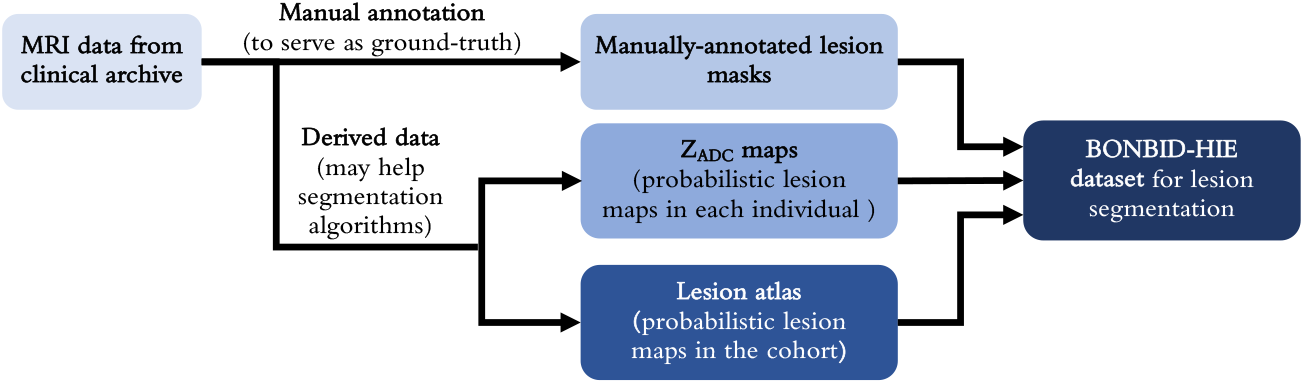
A schematic diagram showing the steps performed on the BONBID-HIE data for release.

### Retrospective Data Collection

Data was retrospectively collected from MGH. Inclusion criteria were: (1) term-born (at physician discretion) (2) clinical diagnosis of HIE; (3) initially treated at MGH between 2001 and 2018; (4) no comorbidities such as hydrocephalus or congenital syndromes; and (5) high-quality MRI acquired in Day 0-14 after birth (visually checked by RW, AF, YO). Exclusion criteria were: (1) excessive motion artifacts or missing images; (2) secondary HIE diagnosis to a primary perinatal stroke.

Clinical characteristics and demographic information were retrospectively gathered from the electronic health records (EHRs). The clinical variables included maternal information during pregnancy and delivery, as well as infant information. More detail can be found in the “Data Records” Section.

MRI data was downloaded from MGH Radiology Department clinical archives using the mi2b2 search engine [29]. MRIs were acquired on either a GE 1.5T Signa scanner (N=52, scanned during 2001-2012), or, a Siemens 3T TrioTim or PrismaFit scanner (N=81, scanned during 2012-2018). Diffusion tensor sequences on all scanners had the protocol as follows: Time of Repetition (TR)=7500 − 9500*ms*, Time of Echo (TE)=80 − 115*ms*, and b=1000 *s/mm*^2^. The GE scanner had resolution × 1.5 × (2.0 − 4.0)*mm*^3^ and (6 - 60) diffusion directions, while the Siemens scanner had a resolution 2 × 2 × 2*mm*^3^ and (25 - 60) diffusion directions. Apparent diffusion coefficient (ADC) maps were directly generated by the scanners (with Syngo software for Siemens scanners [30], and with the Advantage Windows Workstation for GE scanners [31, 32]).

### MRI Pre-processing

Besides the raw NIfTI image as converted from the DICOM files, we also generated several processed images. The pre-processing steps included: N4 bias correction [33], field of view normalization [34], multi-atlas skull stripping for the ADC maps [35], and deformable registration of each patient’s ADC map to a normative 0-14 day neonatal brain ADC atlas [36], by the Deformable Registration via Attribute Matching and Mutual-Saliency weighting (DRAMMS) software [37], which has been extended-validated for lifespan ages in various MRI sequences [38]. This normative ADC atlas was constructed from ADC maps of 13 healthy individuals acquired 0-14 days after birth (Figure 3a) with our extensively-validated MRI analysis pipeline [39, 38, 35, 34]. All software packages used in this pre-processing pipeline are publicly available and have been validated in processing both research and clinical MRI scans across ages [40, 41, 42, 43].

**Figure 3.**
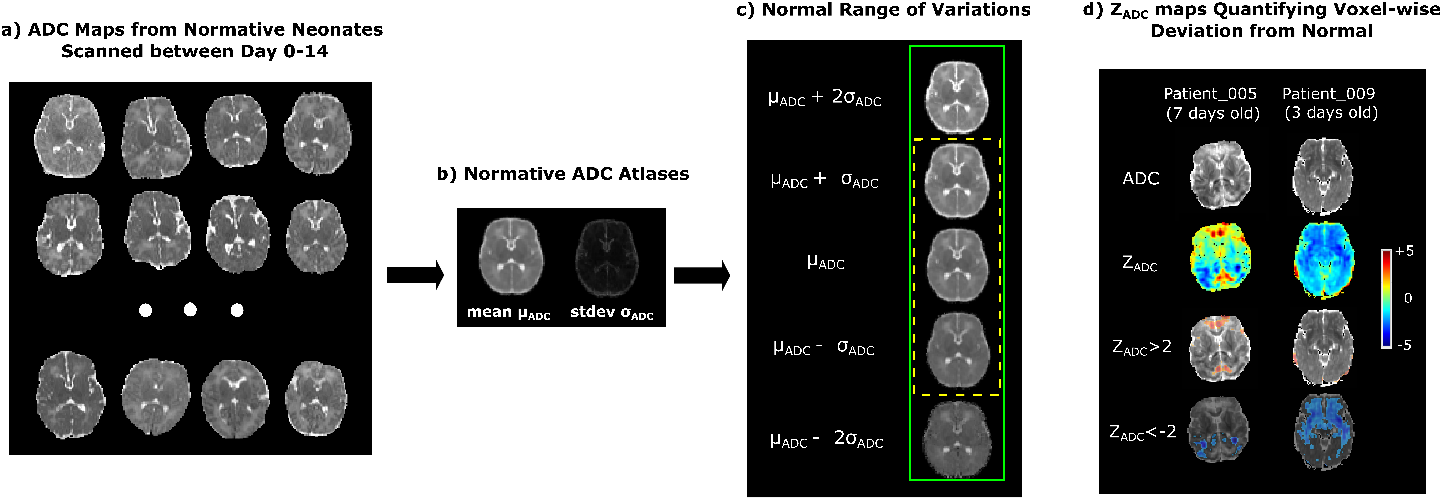
The generation and concept of *Z*_*ADC*_ maps. (a) Examples of ADC maps from normative subjects, which were warped into the same space using unbiased group-wise DRAMMS registration to generate (b) the mean and standard deviation ADC atlases. (c) One or two standard deviations above and below the mean ADC atlas define the normal ranges of voxel-wise ADC variations. (d) Our novel *Z*_*ADC*_ map quantifies voxel-wise deviations from the mean ADC map in (b). The cool/warm colors in (d) represent voxels with ADC values lower/higher than the mean ADC at the same anatomic location, according to the scale bar on the right.

### Expert Annotation of Lesions

HIE lesions were manually annotated as a binary mask on the 3D ADC maps in the patient’s raw image space, using the MRICroN software. ADC maps are clinically used as the standard images to identify HIE abnormalities [44, 45]. The annotations were done by a trained physician (YS; >3 years of experience) according to the neuroradiology reports that were generated as part of the clinical flow. The annotations started from the axial slice and were subsequently modified in the coronal and sagittal planes for the 3D integrity of lesion regions. Uncertainties occurred in 27 patients and were resolved by the consensus of three pediatric neuroradiologists (CJ, SS, and PEG; >5, >5, and >20 years of experience).

### Generation of *Z*_*ADC*_ Maps for Each Patient

Neuroradiologists identify acute brain injury from HIE as regions with low ADC values. Low ADC values represent a reduced water diffusion or blood supply [6]. However, a dilemma is, what ADC value is considered abnormally low versus just low within the normal variation? The normal variations of ADC values differ across brain regions [36, 46], making this question difficult even for experienced neuroradiologists. For example, a voxel with an ADC value of 800 (×10^−6^*mm*^2^*/s*) may be considered normal at one brain region, whereas another voxel with an ADC value of 900 (×10^−6^*mm*^2^*/s*) may be considered lesioned at another brain region, if the normal ranges of ADC variations in the two brain regions are 700-900 and 950-1100 (same unit), respectively.

To address this dilemma, we have developed *Z*_*ADC*_ maps to normalize and make ADC values comparable across brain voxel locations [40]. First, a normative ADC atlas was generated from scans of healthy neonates (Figure 3a). This atlas quantifies the mean ADC values and standard deviation at every voxel [47] (Figure 3b), and hence the normal range of variations at each voxel (Figure 3c). Then, we converted each patient’s ADC map (first row, Figure 3d) into a *Z*_*ADC*_ map (second row, Figure 3d). The *Z*_*ADC*_ maps compared the patient’s ADC value at each voxel to the normal variations at the corresponding voxel in the atlas. In a nutshell, *Z*_*ADC*_ maps quantify how many standard deviations away a patient’s ADC value at a voxel is from the normal mean at the same anatomical location.

Specifically, a deformation (*D*) was computed, which mapped every voxel *x* in the patient’s ADC map to its anatomically-corresponding location *D*(*x*) in the atlas space. The normal range of ADC variation per voxel was defined by the mean and standard deviation denoted for that voxel across all healthy neonates. Finally, the patient’s ADC value *I*_*x*_ at voxel *x* was converted to a *Z* value: *Z*_*x*_ = (*I*_*x*_ *−µD*(*x*))*/σ D*(*x*). We calculated the *Z*_*ADC*_ map, which resides in the patient’s raw ADC image space, for each patient. This offers an option for developing anatomy-aware lesion segmentation algorithms [48].

### Construction of Statistical Lesion Atlases for the Cohort

The same deformation field that was computed by the non-rigid registration from the patient’s skull-stripped ADC map to the normal ADC atlas was used to transform the binary brain lesion maps of each patient into the normal neonatal ADC atlas space [37]. The transformed binary lesion masks were then summed and divided by the total number of patients at each voxel. This led to a statistical lesion atlas that quantifies voxel-wise frequency, or probability, of HIE lesions in our cohort, as illustrated in Figure 4.

**Figure 4.**
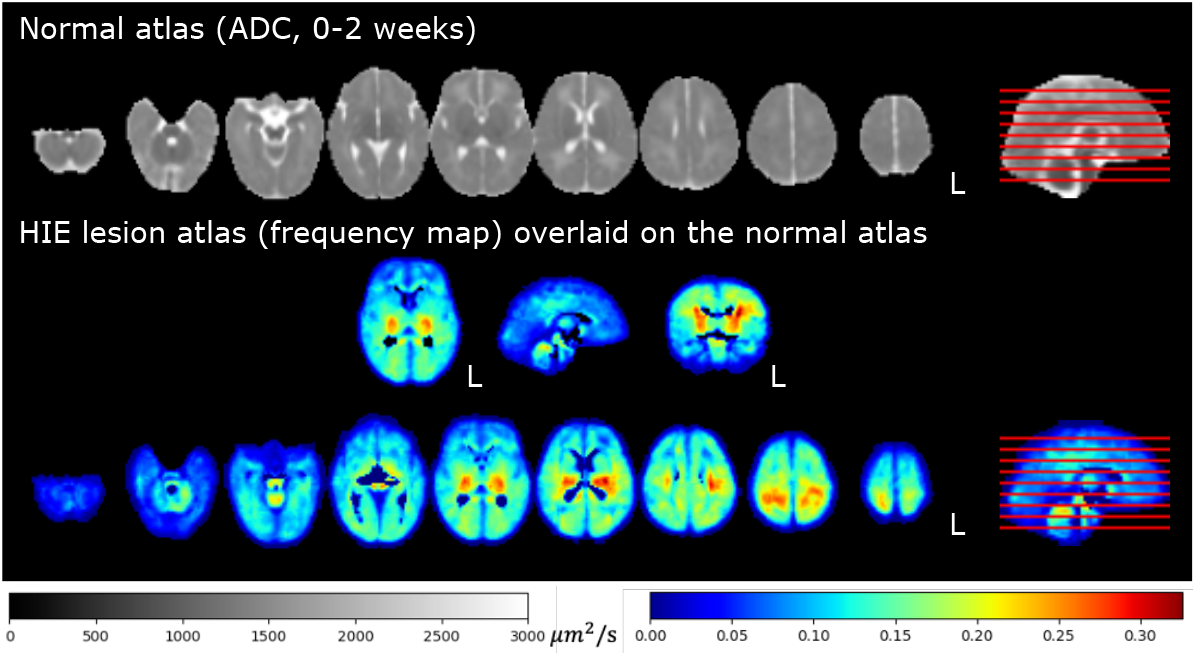
Statistical lesion atlas quantifying the voxel-wise lesion frequency in our cohort of N=133 patients in the normal 0-14 days ADC atlas space.

## Data Records

### Dataset Characteristics

Table 1A lists the demographics and clinical characteristics of mothers and neonates. Maternal information includes demographics (age at delivery, race), birth mode (C-section or vaginal), and complications during pregnancy and delivery. Neonatal information includes demographics (age at MRI scan, gestational age at birth, birth weight, head circumference, sex), birth conditions (1/5/10-minute APGAR scores, lowest pH value in umbilical cord), treatment (hypothermia or not), and complications in the neonatal intensive care unit (NICU), including seizure (yes/no), length of stay (in days), the use of endotracheal tube (ETT, yes/no), and the administration of total parenteral nutrition (TPN, yes/no). In each row, we also listed the number of patients who had such information available.

**Table 1.** Cohort characteristics (N=133)

| **A. Demographics and Clinical Characteristics** |  |  |
| --- | --- | --- |
| **Maternal Information** |  |  |
| Maternal age at delivery (years) | 29.5 ± 6.7 | N=133 |
| Race | White (43), Black or African American (7), Hispanic or Latino (15), Multi Race (5), Unknown (57), Other (6) | N=133 |
| Delivery | C-section (78), Vaginal (55) | N=133 |
| Antepartum hemorrhage | Yes (29), No (104) | N=133 |
| Thyroid dysfunction | Yes (5), No (128) | N=133 |
| Pre-eclampsia | Yes (9), No (124) | N=133 |
| Fetal decels | Yes (72), No (61) | N=133 |
| Shoulder dystocia | Yes (8), No (125) | N=133 |
| Chorioamnionitis | Yes (20), No (108) | N=133 |
| Emergency c-section | Yes (69), No (58) | N=133 |
| **Neonatal Information** |  |  |
| Age at scan (days) | 3.9 ± 2.7 | N=133 |
| Gestational age at birth (weeks) | 39.1 ± 1.9 | N=133 |
| Birth weight (g) | 3321.9 ± 615.9 | N=133 |
| Infant head circumference (cm) | 34.2 ± 1.4 | N=85 |
| Sex | Male (74), Female (59) | N=133 |
| 1-minute APGAR scores | 1.9 ± 1.7 | N=133 |
| 5-minute APGAR scores | 4.2 ± 2.3 | N=132 |
| 10-minute APGAR scores | 5.3 ± 2.1 | N=118 |
| Lowest pH value in umbilical cord | 7.00 ± 0.20 | N=129 |
| Therapeutic hypothermia before MRI? | Yes (86), No (47) | N=133 |
| Endotracheal tube (ETT) in NICU | Yes (78), No (47) | N=125 |
| Total parenteral nutrition (TPN) in NICU | Yes (111), No (21) | N=133 |
| Seizures NICU | Yes (64), No (69) | N=133 |
| Length of stay in NICU (days) | 12.10 ± 9.92 | N=133 |
| **B. Lesion Characteristics (N=133)** |  |  |
| Whole-brain lesion volume – minimum | 0 mm<sup>3</sup> (0% of the brain injured) |  |
| Whole-brain lesion volume – 25th percentile | 441.96 mm<sup>3</sup> (0.10% of the brain injured) |  |
| Whole-brain lesion volume – median | 2765.63 mm<sup>3</sup> (0.63% of the brain injured) |  |
| Whole-brain lesion volume – 75th percentile | 24264.27 mm<sup>3</sup> (4.86% of the brain injured) |  |
| Whole-brain lesion volume – maximum | 412120.00 mm<sup>3</sup> (82.59% of the brain injured) |  |
| **C. Number of Patients by Percentage of Lesions in the Brain (N=133)** |  |  |
| [0%, 1%) of the brain being injured | 55.64% (N=74) |  |
| [1, 5%) of the brain was injured | 19.55% (N=26) |  |
| [5, 10%) of the brain was injured | 5.26% (N=7) |  |
| [10, 20%) of the brain was injured | 4.51% (N=6) |  |
| [20, 50%) of the brain was injured | 8.27% (N=11) |  |
| [50, 100%] of the brain was injured | 6.77% (N=9) |  |
| **D. Scanner (N=133)** |  |  |
| GE 1.5T | 39% (N=52) |  |
| Siemens 3T | 61% (N=81) |  |

Table 1B quantifies the distribution of the absolute lesion volumes (in *mm*^3^) and relative lesion volume (percentage of the brain being injured). Here, the relative lesion volume was calculated by the volume of the expert-annotated lesion regions divided by the algorithm-extracted whole brain volumes [34, 35]. The median lesion volume accounted for 0.63% of the whole brain volume. This confirms that over half of the patients had less than 1% of the brain being injured. Table 1C further calculates the distribution of the relative lesion volumes (by the percentage of the brain being lesioned). The absolute and relative volumes were both computed in the patient’s raw ADC image space. The minimum lesion was 0 *mm*^3^, which is a common issue in HIE – mild HIE cases may not show explicit lesions in neonatal MRIs [49, 50, 51].

Figure 5 shows the ADC map, *Z*_*ADC*_ map, and expert annotations of example patients with different HIE lesion percentages. Two patients are shown in each of the four groups: those with lesions occupying <1% (upper left panel), 1-5% (upper right panel), 5-50% (lower left panel), and 50-100% (lower right panel) of the whole brain volume. Overall, around 1 in 2 (55.64%) patients had HIE lesions occupying less than 1% of their brain volume, and 3 in 4 patients (75.19%) patients had lesions occupying less than 5% of their brain. This confirms that HIE lesions detectable in the diffusion MRI in our cohort are often small.

**Figure 5.**
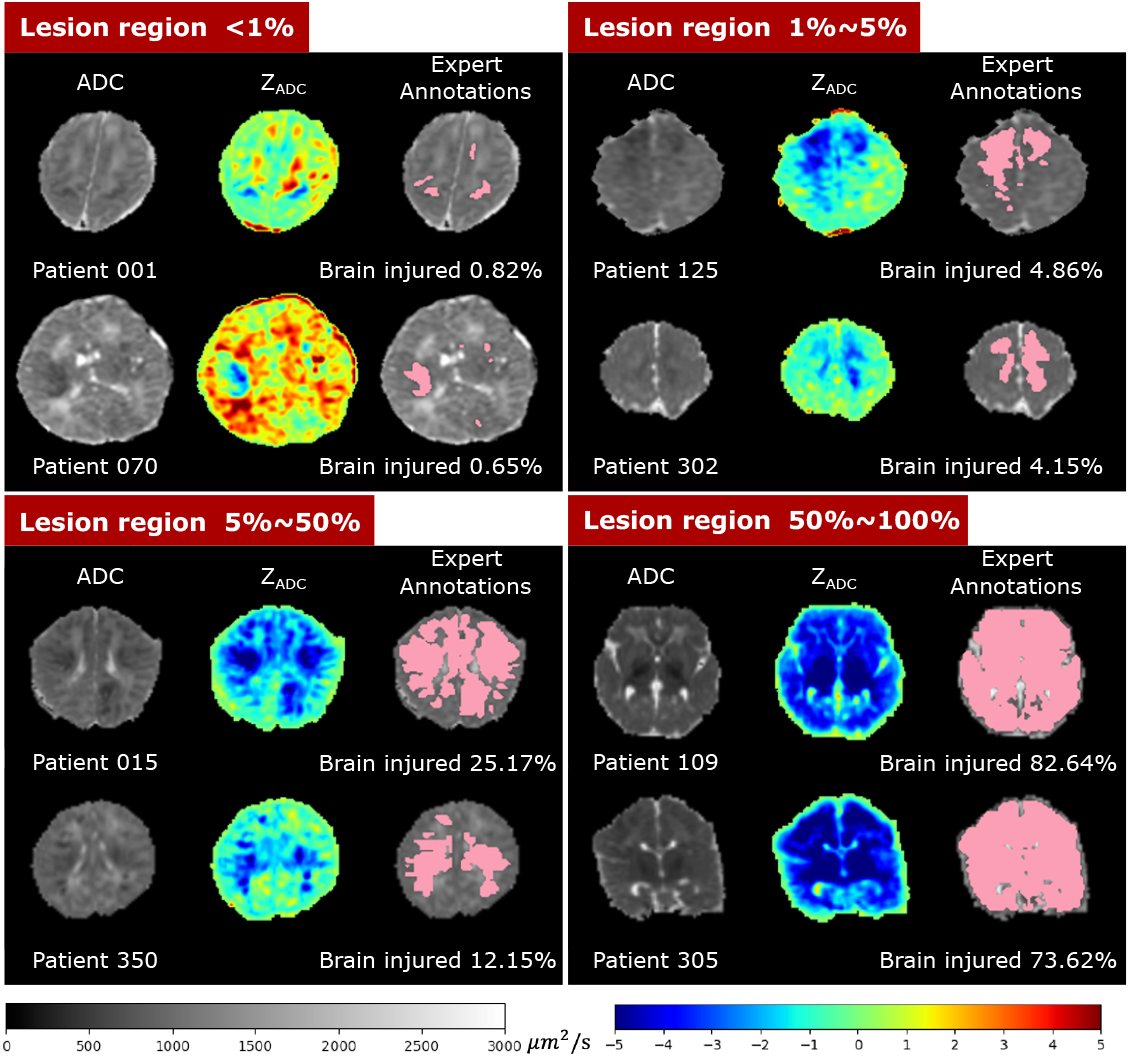
Visualization of patients with different lesion percentages. In every patient, the left image is the ADC map (skull stripped) with range of ADC values designated by the gray scale bar, the middle is the computed *Z*_*ADC*_ map with range of Z-scores designated by the rainbow scale bar, and the right image is the expert-annotated lesion regions (pink) overlaid on the ADC map. Percentages of injury were calculated by the volume of the expert-annotated lesion regions divided by algorithm-extracted whole brain volumes.

### Data structure and file formats

All medical imaging files were exported from the Picture Archiving and Communication System (PACS) and converted into the NIfTI format. Segmentation masks created by expert annotations were also saved in NIfTI format. Corresponding scanner metadata from the Digital Imaging and Communications in Medicine (DICOM) header in the .json file format is provided with the datasets. All data in the BONBID-HIE dataset was separated into a training dataset (n=89) and a test dataset (n=44). Both the training and test sets contain data from both scanners (GE 1.5T Signa and Siemens 3T Trio). The split between the training and test data set has been performed (RB, YO) so that both sets include a similar variance of HIE lesion patterns as shown in Table 1C.

The data is organized in the format shown in Figure 6. BONBID-HIE provides, per patient: (i) 0ADC: raw defaced ADC maps; (ii) 1ADC_ss: skull stripped ADC map; (iii) 2Z_ADC: *Z*_*ADC*_ map; (iv) 3LABEL: expert lesion annotations; and (v) clinical data: clinical variables as written in Table 1A. There is also (vi) Atlases: a folder for the normal and lesion atlases; (vii) a readme.txt file: a text file to provide information on this data organization; and (viii) the license file of the BONBID-HIE dataset.

**Figure 6.**
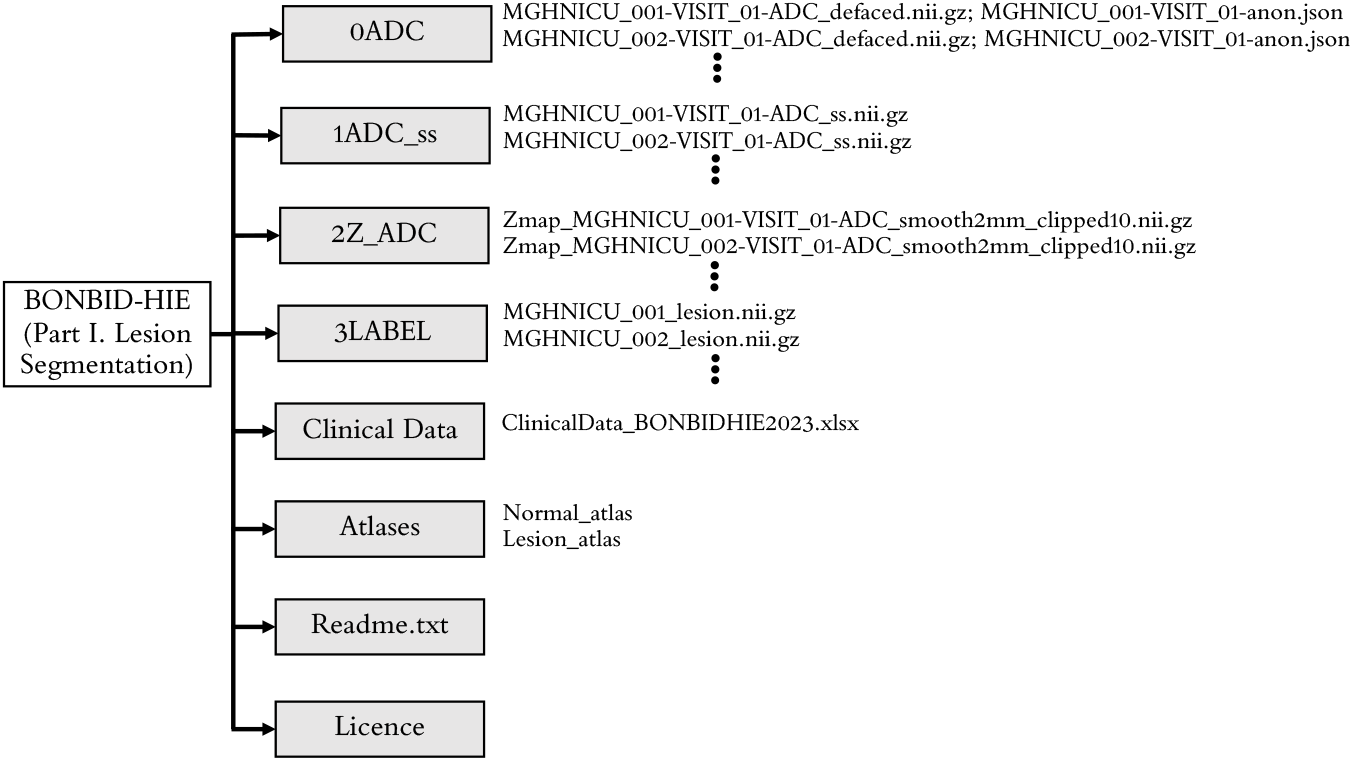
Folder structure of the BONBID-HIE dataset (Part I. Lesion Segmentation).

### Data repository and storage

All training data has been made publicly available under the CC BY NC ND license (https://creativecommons.org/licenses/by-nc-nd/2.0/, allowing academic use with credit, prohibiting commercial use without owner’s permission, and disallowing derivation or adaption of data). This dataset is also used for the HIE lesion segmentation challenge in Medical Image Computing and Computer Assisted Intervention (MICCAI) 2023 annual conference in October 2023. Further information about the HIE challenges, including data storage and download, can be found under our homepage (https://hiechallenge.github.io/). As such, the expert lesion annotation of lesions in the testing sub-cohort will be released after the challenge.

### Technical Validation

#### Representativeness of patient cohort

Our data is representative of HIE cohorts in the developed countries. At least three characteristics of our data agree with documented clinical knowledge about HIE.

##### Lesion distribution in space agrees with clinical knowledge

Our statistical lesion atlas in Figure 4 shows that HIE lesions can occur anywhere in the brain. The regions most frequently injured here included the basal ganglia, internal capsules, thalamus, temporal lobes, cerebral white matter, brainstem, and cerebellum (red, orange, and yellow regions in Figure 4). This lesion atlas map coincides with clinical knowledge of brain regions often vulnerable to HIE injuries [2, 52, 6]. Indeed, HIE-related injuries in these regions have been key criteria in expert MRI scoring systems, which are used to assess the severity of HIE. Examples include the NICHD Neonatal Research Network (NRN) [2], the Barkovich [53], the Weeke-deVeries [3], and the Trivedi [54] scoring systems. In addition, lesions appeared in less than 35% of the patients at any given voxel, according to the color bar in this figure. This confirms the clinical knowledge that HIE lesions are diffuse, spatially distributed, and almost half to two-thirds of the HIE patients are mild to moderate, at least in patients in the USA [50, 51].

##### Lesion distribution in time agrees with clinical knowledge

Figure 7(a) shows the percentage of the whole brain volume being lesioned at different postnatal ages. The lesion percentage in the ADC maps came down to almost 0 in the 9 patients who underwent MRI scans after postnatal day 7. This agrees with the clinical knowledge that HIE-related lesions are more detectable in ADC during 0-7 postnatal days or in T1/T2-weighted images than in relatively later scans (after postnatal day 7) [55, 51, 2].

**Figure 7.**
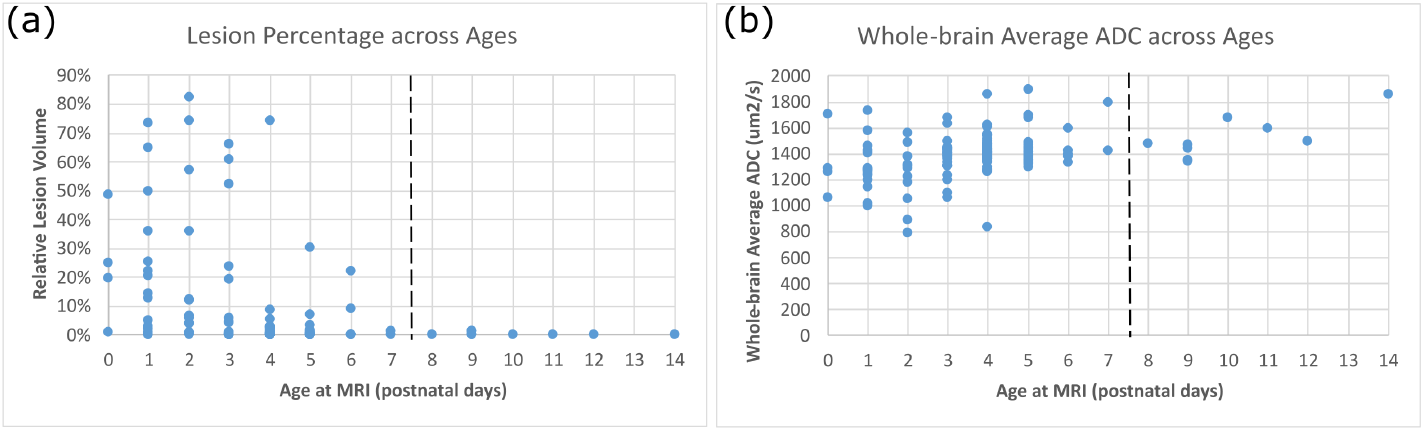
Representativeness of our cohort for (a) lesion distribution across ages; and (b) whole-brain ADC values across ages. In both panels, each dot denotes a patient in our cohort.

##### ADC evolvement with age agrees with clinical knowledge

Figure 7(b) shows the whole-brain average ADC values of all patients (each dot is a patient). In normal cohorts, ADC values drop rapidly in the early postnatal life (see Figures 4 and 5 in [36], and Figures 3 and 4 in [46]). However, the presence of HIE-related abnormalities disrupted this trend – HIE patients undergoing earlier MRIs (0-7 postnatal days) had decreased ADC values in a larger precentage of the brain (Figure 7(a)), so the ADC values in 0-7 postnatal days were at similar or even lower levels than ADC values in 7-14 postnatal days among HIE patients (Figure 7(b)). This has also been documented in HIE literature [56, 57].

#### Utility of ADC maps and *Z*_*ADC*_ maps

To demonstrate the utility of the computed *Z*_*ADC*_ maps, we compared the accuracies of using ADC or *Z*_*ADC*_ maps for lesion segmentation. We attempted simple thresholding of ADC and *Z*_*ADC*_ maps, at several threshold values, for segmenting HIE lesions. Although simple, thresholding-based segmentation accuracy is a strong indicator for segmentation accuracies in more sophisticated machine/deep learning algorithms [58]. For ADC maps, we used different thresholds ranging from 800-1100 *µm*^2^*/s*, as suggested in the literature [55, 59, 60, 61, 62]. For *Z*_*ADC*_ maps, we used thresholds -1.5, -2, and -2.5. Voxels in patient MRIs with ADC values 1.5 to 2.5 standard deviations below the average ADC values from healthy controls were considered abnormally low and hence, lesioned. The choice of thresholds around -2 in *Z*_*ADC*_ maps was also based on the normal distribution of ADC values at each voxel across subjects [36, 40].

We evaluated the accuracy of these maps compared to expert-annotated ground-truth masks using the Dice coefficient, sensitivity, and specificity. Results are shown in Figure 8. Here, the gray boxplots are the accuracy measurements when ADC maps were thresholded between 800 and 1100 *µm*^2^*/s* (50 *µm*^2^*/s* intervals). The blue boxplots are the accuracy measurements when the *Z*_*ADC*_ maps were thresholded at -1.5, -2, and -2.5. Figure 8 demonstrates: (i) both ADC and *Z*_*ADC*_ have value in helping segment the HIE-related lesions, since the specificity from simple naive thresholding-based segmentations was comparable to those from machine learning-based algorithms [20], although the Dice and sensitivity were lower; (ii) *Z*_*ADC*_ maps thresholded at -2, the most intuitive and straightforward threshold value, yielded the highest Dice (0.54 ± 0.28), followed by *Z*_*ADC*_ maps thresholded at -2.5 (Dice 0.39 ± 0.25); (iii) *Z*_*ADC*_ maps thresholded at -1.5 yielded the highest 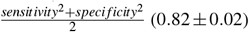 compared with any ADC thresholds, followed by *Z*_*ADC*_ maps thresholded at -2 (0.76 ± 0.03); and (iv) overall, across all thresholds, *Z*_*ADC*_ maps showed a higher area under the curve (AUC: 0.936). This shows that *Z*_*ADC*_ maps – anatomy-normalized ADC images – carry the potential to improve lesion detection accuracy.

**Figure 8.**
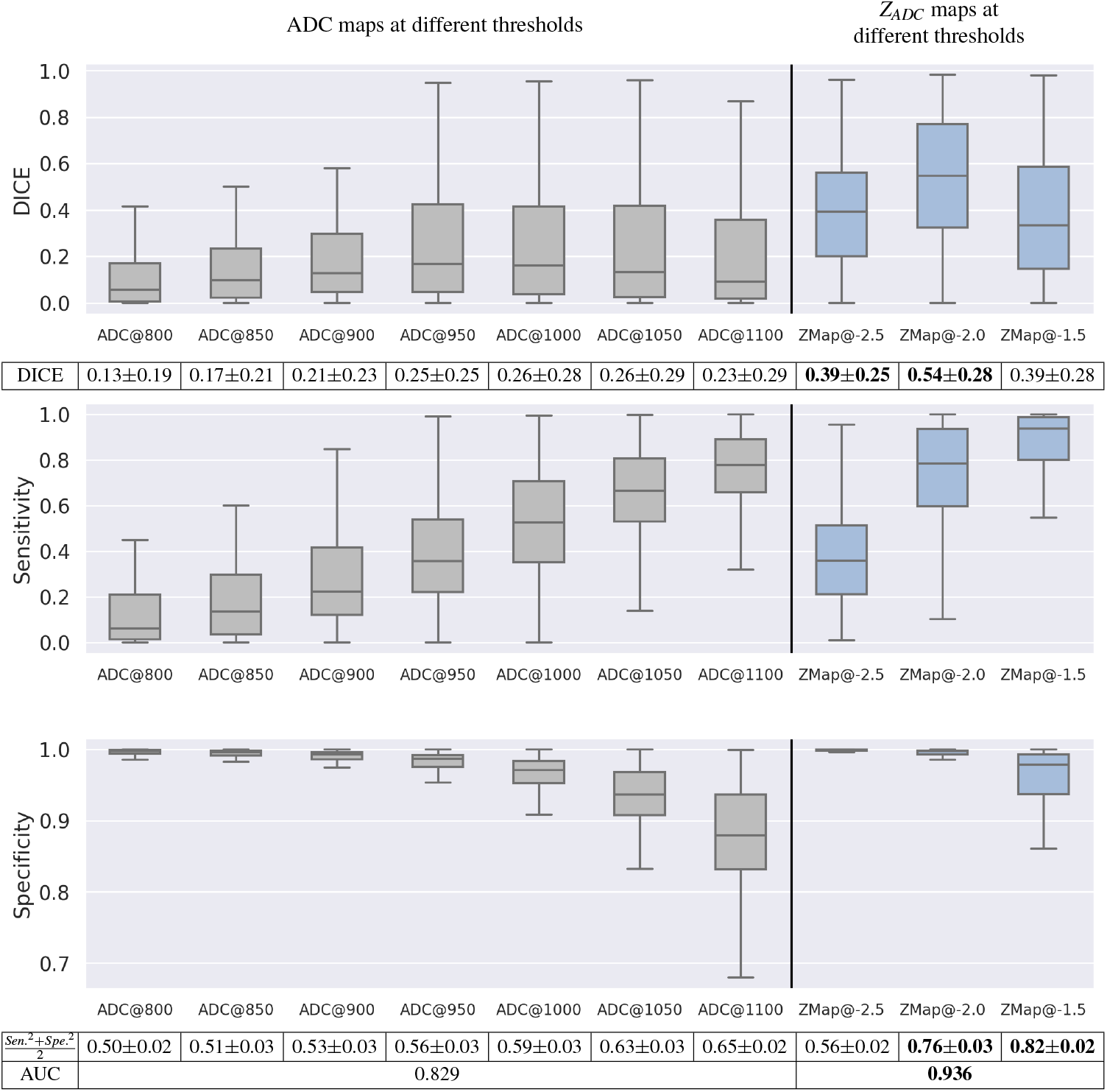
Accuracy of thresholding-based lesion segmentation on ADC and *Z*_*ADC*_ maps with different threshold values. Bold texts in the tables beneath the figure panels highlight the two scenarios with the highest Dice scores, the two scenarios with the highest and most balanced sensitivity and specificity metrics, and the parametric map (ADC or *Z*_*ADC*_) with the highest area under the receiver-operating-characteristic curve (AUC).

## Code Availability

Data is available as part of the 1^*st*^ BONBID-HIE MICCAI challenge (https://hiechallenge.github.io/), to be held in October 2023. For the training sub-cohort, all MRI (raw and derived) and expert lesions annotations are available. For the testing sub-cohort, only the MRI (raw and derived) data is available at the time this manuscript is drafted, and the expert lesion annotations will be made available after the challenge in October 2023.

Codes to automatically calculate the evaluation metrics are available in BONBID-HIE 2023 MICCAI challenge (https://bonbid-hie2023.grand-challenge.org/). The repository in the challenge contains scripts to read the images, visualize them, and quantify the algorithm’s performance with the same metrics used in the challenge to rank participants.

## Acknowledgements

This work was supported, in part, by NIH R03HD104891, R21NS121735, and R61NS126792.

